# Blender tissue cartography: an intuitive tool for the analysis of dynamic 3D microscopy data

**DOI:** 10.1101/2025.02.04.636523

**Authors:** Nikolas Claussen, Cécile Regis, Susan Wopat, Matthew F. Lefebvre, Sebastian Streichan

## Abstract

Volumetric microscopy can image complex 3D tissues, but 3D image data remains difficult to visualize and quantify. Many biological systems are organized as thin, curved sheets (for example, epithelia). Tissue cartography extracts and cartographically projects these curved surfaces from volumetric images. This converts 3D into 2D image data, greatly facilitating visualization, analysis, and computational processing. Existing tools, however, demand advanced coding expertise and are limited to simple tissue geometries. Here, we present `blender issue cartography` (`btc`), an interactive add-on for the 3D editor Blender that makes tissue cartography user-friendly by a graphical interface, and handles complex biological shapes using powerful computer graphics algorithms. An accompanying Python library supports faithful 3D measurements in 2D cartographic projections and custom analysis pipelines. Time-lapse data can be batch-processed by algorithmically aligning all time points to a single *key frame*. We demonstrate `btc` on diverse and complex tissue shapes from *Drosophila*, stem-cell organoids, *Arabidopsis*, and zebrafish. `btc` enables quantitative cartographic analysis of complex 3D tissues, broadening access to methods previously restricted to specialists, while leveraging tools from computer graphics to unlock new capabilities.

## Main

Biological tissues, organs, and whole embryos are inherently three-dimensional (3D), requiring volumetric microscopy, yet 3D image data is difficult to store, visualize, and analyze quantitatively. Tissue cartography exploits the layered organization of epithelia, leaves, or tube-like visceral organs to represent such structures as curved 2D surfaces embedded in 3D[1]. Workflows segment out a *surface of interest* (SOI) from volumetric data (e.g., confocal *z*-stacks) and *unwrap* the SOI to the 2D plane, enabling 2D visualization and analysis of 3D data. This facilitates cell segmentation and tracking [2], quantification of tissue deformation [3] and protein localization patterns [4], or *in-toto* visualization of curved objects. These methods have notably proven useful in developmental biology, where they can extract salient biological features, like tissue layers and dynamic organ geometry. Cartography can also register recordings across samples or time to a common reference [5]. Tissue cartography aims to advance quantitative understanding of development in the spirit of D’Arcy Thompson [6], whose vision of development as a series of “cartographic transformations” continues to inspire current research [7].

Existing tools for tissue cartography have enabled striking, quantitative insights into 3D morphogenesis [8, 2, 9]. However, limitations impede widespread adoption and analysis of complex, dynamic geometries. Tools like `LocalZProjector` [10] parameterize surfaces via a *height function* and hence cannot handle fully 3D surfaces with “overhang”. The powerful 3D analysis software `MorphoGraphX` [11] has enabled sophisticated 3D analyses in plant biology. However, `MorphoGraphX` does not perform cartographic projections; instead, it projects image intensity onto the vertices of a high-resolution mesh. However, in 3D, only part of a curved surface is visible at any time. Further, the resulting mesh-associated data is more complex and less flexible than cartographic projections, and less compatible with machine-learning-based image analysis software. These issues are exacerbated if the image is difficult to segment into cells. The `ImSAnE` [1] and `TubULAR` [3] software package can carry out tissue cartography for more complex shapes. However, `TubULAR` is specialized to tube-like surfaces only. Further, in both packages, reliance on specialized MATLAB code and the lack of interactive, graphical surface/projection editing hinder non-expert use. Lastly, `ImSAnE` predates many modern surface-processing algorithms [12]. Here, we present `blender tissue cartography` (`btc`), an add-on to the popular, open-source 3D editor Blender [13] and an accompanying Python library. `btc` makes tissue cartography of dynamic, highly curved surfaces accessible through a unified, modular pipeline with a graphical user interface, leveraging high-quality computer graphics and 3D-animation tools.

## Results

### 1 Tissue cartography workflow in btc

Tissue cartography generally proceeds in the following steps (Fig. 1A-A”):

**Figure 1:**
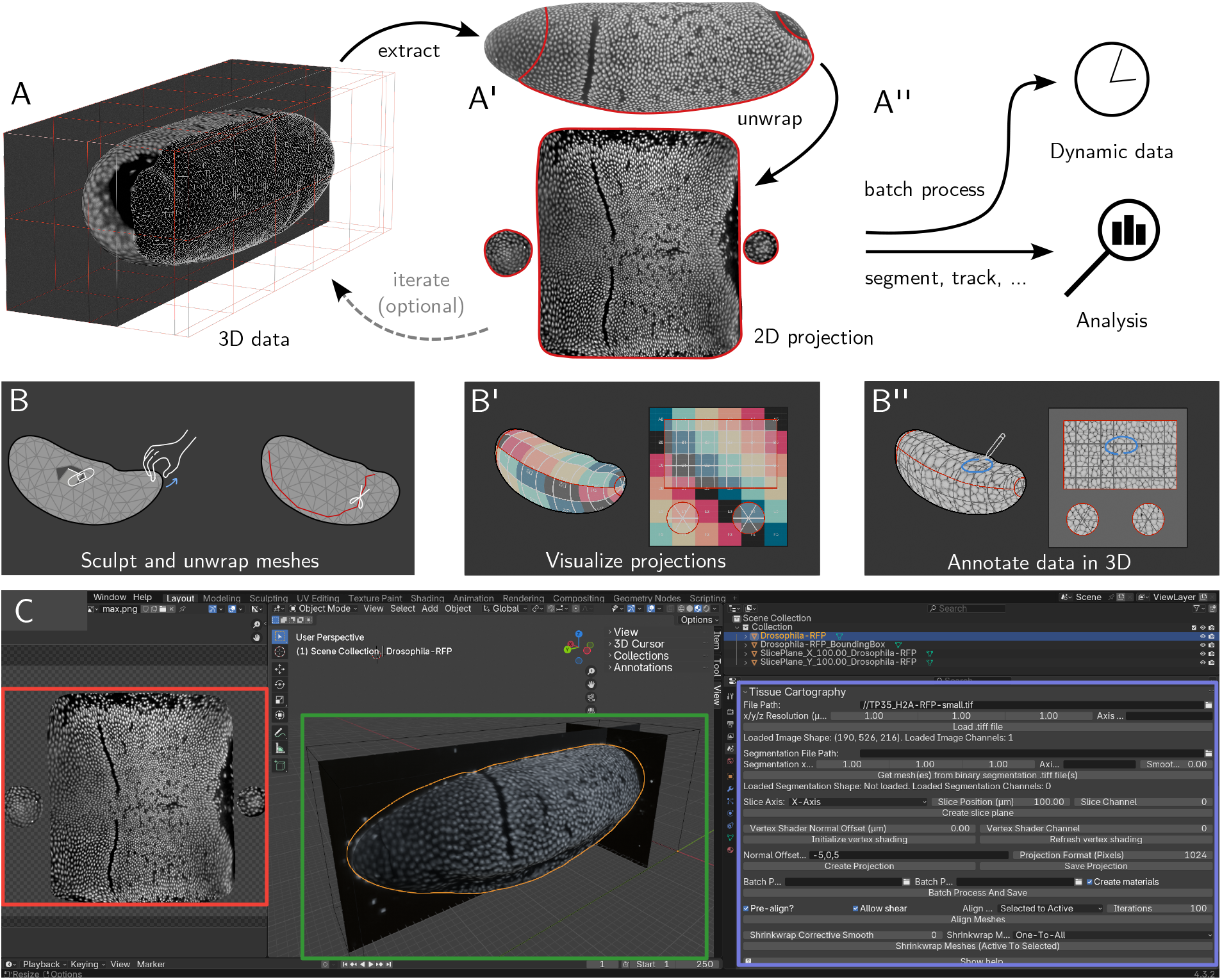
btc for user-friendly 3D analysis workflows. **A-A”** (A) 3D image data image (*Drosophila* blastoderm with fluorescently labeled nuclei, anterior left, ventral down). (A’) The extracted curved 2D surface of interest is represented by a polygonal mesh. Surface unwrapping (UV mapping) projects 3D image data into 2D. After inspection, meshes and UV maps can be rapidly edited to improve the quality of 2D projections. (A”) 2D projections can be used for *in toto* visualization or downstream analyses like segmentation. Dynamic datasets can be batch-processed (see Fig. 3). Modular design allows switching out the tool used in each pipeline step. **B-B”** Interactive 3D data processing in Blender’s GUI. (B) Surface sculpting to fix segmentation errors. (B’) Blender’s UV editor combines powerful cartographic algorithms with graphical user input (e.g., placement of cartographic cuts). Cartographic projections can be visualized in 3D. (B”) Projected data can be visualized and annotated in 3D. **C** Blender user interface. Cartographic projection (red), surface of interest with 3D data visualized by orthoslices and projected mesh intensity (green), `btc` add-on toolset for loading and processing 3D data (purple).

1. **Extraction:** Detect the surface of interest in volumetric image data, typically via 3D image *segmentation*, and convert the result to a surface mesh (*meshing*).
2. **Unwrapping:** Unwrap the 3D surface to a 2D cartographic plane.
3. **Image projection:** Project 3D image data into 2D.
4. **Batch processing:** Process all frames of a time-lapse recording.
5. **Analysis and visualization:** Measure quantities of interest in the 2D projection, accounting for the surface’s 3D curvature.

We now describe how these steps are implemented in `btc`. `btc` comprises two tools:

1. **The** `btc` **Blender add-on** carries out tissue cartography within a graphical user interface without requiring any coding (Fig. 1C). The add-on makes Blender’s 3D toolset (surface sculpting, unwrapping, and annotation, Fig. 1B-B”) available for the analyses of 3D image datasets. 3D data can be visualized throughout (facilitating, for instance, choosing a suitable cartographic projection).
2. **The** `btc` **Python library** is an open-source Python package available on pip and allows advanced users to create custom analysis pipelines. Each pipeline step terminates in a single file of standardized type (e.g., a `.obj` mesh file). The modular design allows switching out the tool used for each step. Tutorial Jupyter notebooks can serve as starting points for specialized pipelines.

Comprehensive documentation for the library and add-on, tutorials, a set of template analysis pipelines, and a 2-minute video demonstration can be found online.

### Extraction: segmentation and meshing

To extract the surface of interest, the volumetric image is first segmented in 3D, detecting either (a) the voxels that are part of the SOI or (b) the voxels that are part of the solid object whose boundary is the SOI (see examples in Fig. 4A-A”). `btc` provides an interface for binary segmentation using the popular machine learning software `ilastik` [14] and via level set algorithms [15]. Next, the resulting binary mask is converted into a surface mesh. Throughout `btc`, surfaces are represented as polygonal meshes in the standard `.obj` format (see SI A). `btc` contains a variety of functions for mesh quality improvement (remeshing), smoothing, and processing. Within `Blender`, the resulting meshes can be inspected. Potential segmentation errors can be fixed using `Blender`’s 3D editing tools (Fig. 1B). `btc` also makes surface reconstruction tools from `MeshLab` [16] available in Blender to streamline the workflow. The volumetric image data can also be loaded into `Blender`, allowing the user to sculpt the mesh while visualizing the image data in 3D or projected onto the mesh surface.

### Surface unwrapping and image projection

Now, the mesh can be cartographically unwrapped to a 2D plane (Fig. 1A’). This is known as *UV mapping* in the graphics community. `Blender` possesses a powerful graphical UV editor that implements several cartographic algorithms, from standard axial, spherical, and cylindrical projections to powerful, state-of-the-art algorithms like SLIM [12] which can unwrap even complex surfaces with minimal distortion. Additionally, `Blender` allows the user to define the location of *seams* (cartographic cuts, (Fig. 1B), manually fine-tune the results, and visualize the projection (Fig. 1B’). The resulting UV map defines how the 3D surface is mapped to the plane. The mesh and its UV map can be exported to or loaded from a standard `.obj` file. `btc` then uses an interpolation algorithm to project voxel intensities from the volumetric data onto the 2D cartographic plane. We refer to the result as a *2D projection* (Fig. 1A’, bottom). `btc` also features a second, rapid shading algorithm to visualize image intensities on the mesh surface without a UV map, for example, while sculpting or unwrapping it. In Blender, the intersection of SOI and 3D image data is visualized live, so the user can easily check which part of the 3D data is being extracted.

### Multilayer projections

`btc` can create multi-layer projections, analogous to peeling off the layers of an onion. Each layer is offset inwards/outwards from the SOI along the surface normals. Fig. 2 shows multilayer projections of an *in toto* recording of 12hpf (hours post fertilization) zebrafish embryo. The embryo is unwrapped using a spherical projection (Figs. 2A-B). The user can visualize the image intensity while rotating the sphere to align the equator of the spherical projection with the anterior-posterior axis of the animal. Figs. 2C-C’ shows two-layer projections, at the surface of the embryo and 20*μ*m inwards, where the mesoderm and forming somites are visible.

**Figure 2:**
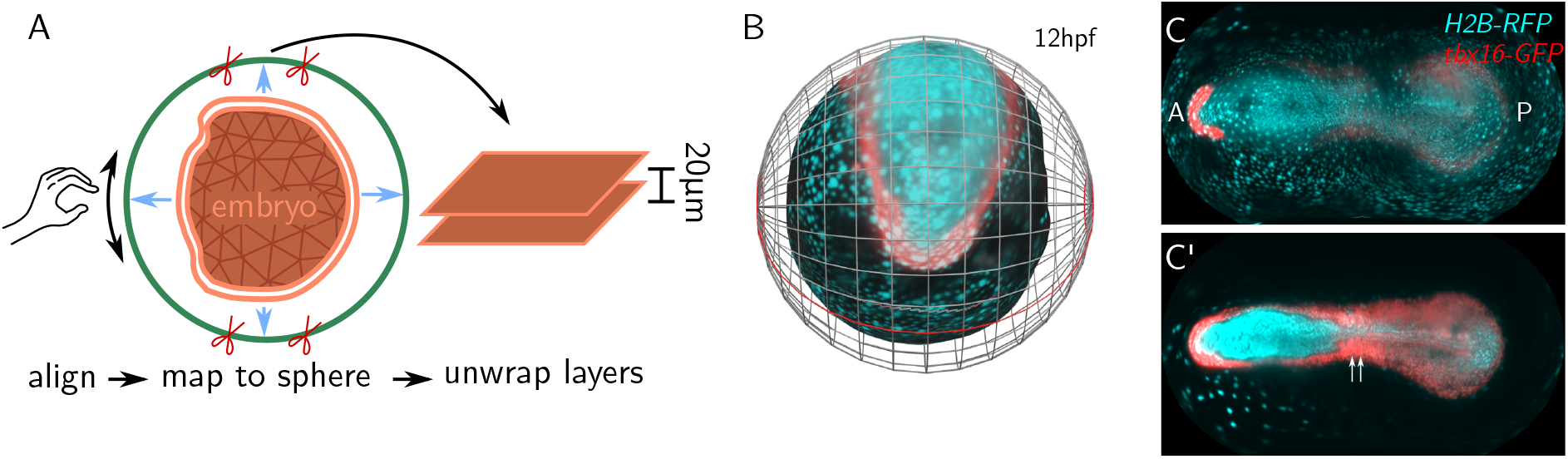
Extraction of multiple tissue layers. **A** Zebrafish embryo (orange mesh) is segmented from an *in toto* lightsheet recording. The embryo’s surface is projected to a sphere, which is then unwrapped. Rotation of the sphere allows visual alignment of the cartographic projection with the AP axis. `btc` extracts multiple tissue layers in a common cartographic projection. **B** Surface of a zebrafish embryo at 12hpf (neurolation) with sphere used for cartographic projection. Fluorescent intensities of 3D data are projected onto the embryo surface (cyan: H2B-RFP (nuclei), red: tbx16-GFP [17] (mesoderm)). Cartographic cut highlighted in red. **C-C’** Fluorescent intensities cartographically projected into 2D, at the surface of the embryo (C) and 20*μ*m below (C’). Mesoderm (tbx16-positive) is localized in the deeper layer forms somites (white arrows). The anterior tbx16-positive region is the prechordal plate.

### Analysis and visualization

`btc` saves the 2D projection (and 3D positions for each pixel in the projection) as `.tif` stacks for 2D visualization and analysis, and as textures for visualization on the 3D mesh. This allows rendering high-quality figures and rapid, iterative improvements of the mesh and its UV map. For instance, the user can adjust the placement of cartographic seams based on the image textures, e.g., focusing on a region of particular interest. Further, `Blender` can annotate data in 3D, with results (e.g. cell labels) saved on top of 2D projections (Fig. 1B”).

For quantitative analysis, `btc` provides a suite for measuring and correcting for cartographic distortion and mapping quantities measured in 2D back (like cell tracks or outlines) back to 3D. For instance, these tools can correctly compute the area of a cell in a 2D projection, even if the unwrapping introduces local area distortion (Fig. S1A-B). We also provide tools to analyze vectorial fields, like morphogenetic tissue flows (SI B).

### 2 Batch processing and dynamic datasets

Dynamic datasets (movies) are represented in `btc` frame-by-frame (one surface mesh and volumetric image file per timepoint). The `btc` add-on or Python library can then *batch-process* all frames of a movie. Users can also batch-process multiple recordings, for example, different replicates of the same experiment. In `btc`, the user defines a UV map for a selected *reference timepoint* – e.g., the first or last frame of a movie. The reference is then algorithmically mapped onto the meshes of the remaining time points via *surface-to-surface registration* algorithms (Fig. 3A-B and SI C). This *transfers* the UV map from the reference to all other timepoints or samples, creating coherent 2D projections in a *common reference frame*. When batch-processing multiple samples, surface-to-surface alignment allows combining recordings of, say, different markers into a single “atlas” [5]. Surface-to-surface alignment is a key step in quantitative and comparative analysis of 3D image data across experimental conditions. We validated `btc`’s approach to time-lapse data on an example of a highly dynamic surface, the embryonic *Drosophila* midgut (Fig. 3C-D, data from Ref. [2]).

**Figure 3:**
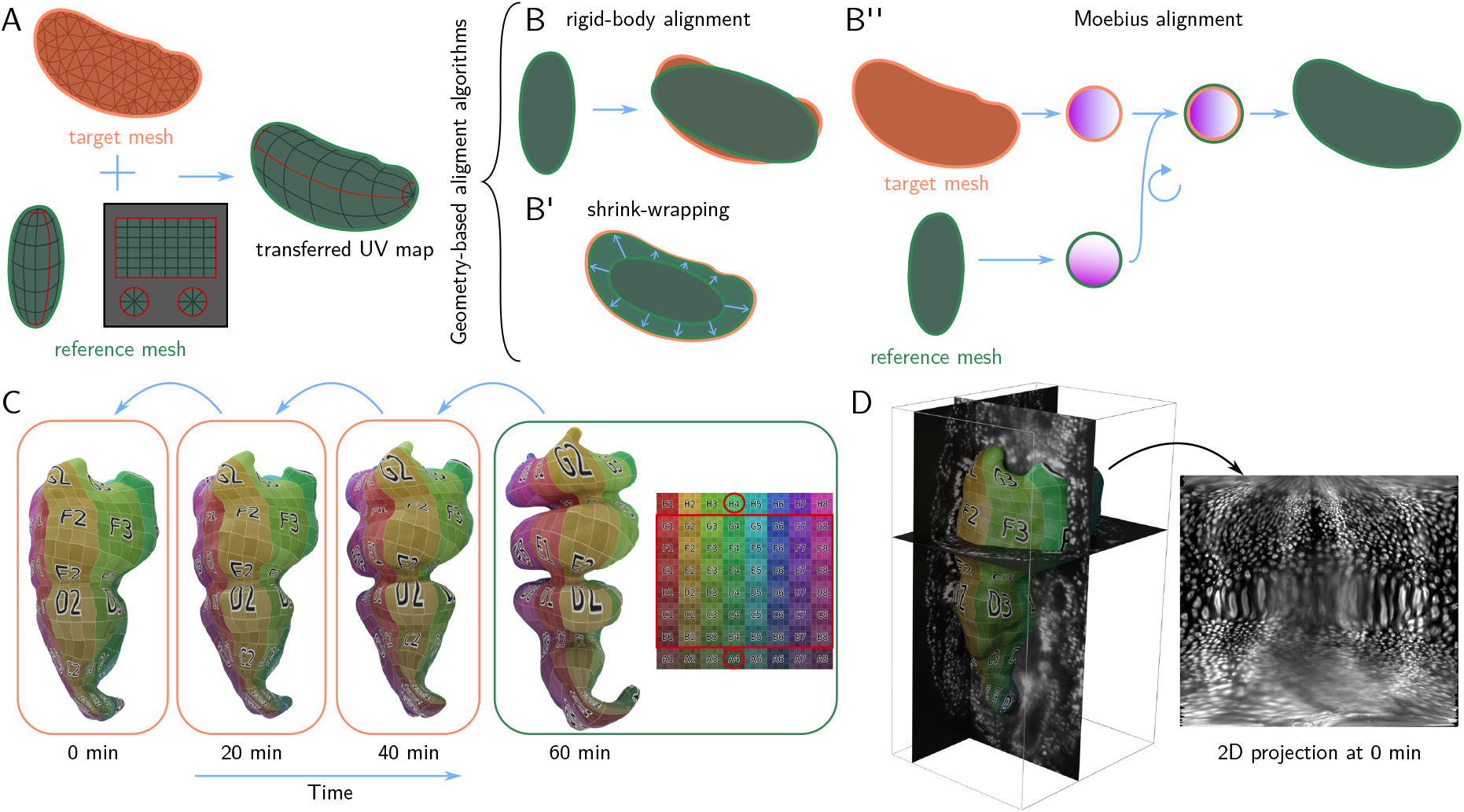
Batch-processing and dynamic data via UV surface-surface alignment. **A** Transferring a UV map to the orange target mesh from the green reference mesh via surfaceto-surface alignment: the reference mesh is deformed to match the shape of the target mesh while preserving its UV map. **B-B”** Surface-to-surface alignment algorithms: rigid-body alignment, shrink-wrapping (smoothed closest-point projection), and Moebius alignment (see text). All algorithms are based on the shape of the surface only. **C** UV transfer for a highly dynamic biological surface. Mesh of the embryonic *Drosophila* midgut extracted from live deep-tissue *in-toto* imaging (data from [2]). The complex shape of the final timepoint is unwrapped in `Blender` and the UV map is iteratively transferred to all other timepoints via shrink-wrapping. **D** The extracted surface at timepoint 0 together with 3D data (fluorescently marker nuclei) and 2D projection.

**Figure 4:**
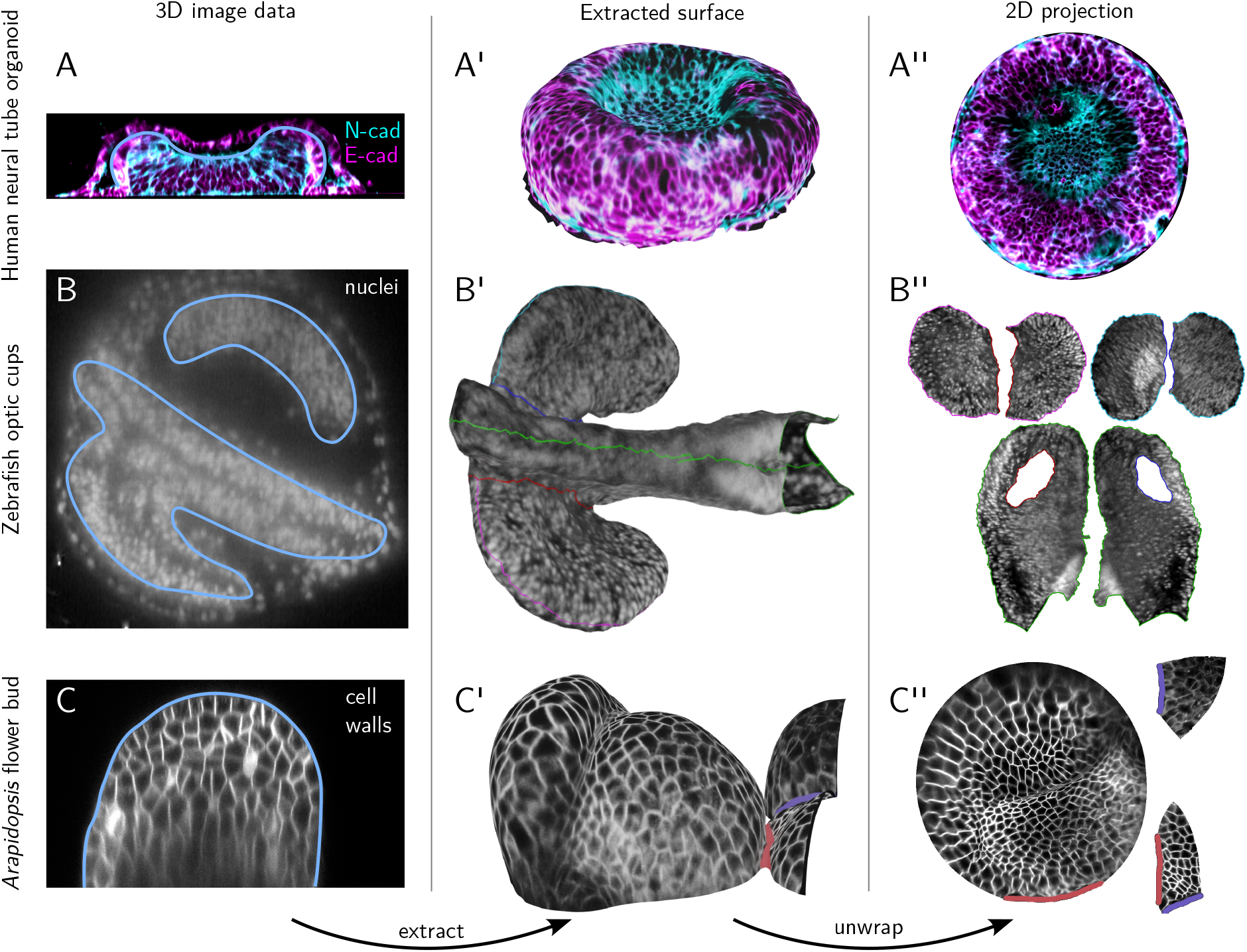
btc processes complex and diverse biological shapes. **A-A”** Confocal *z*stack of a human neural tube organoid [18], stained for neural and epithelial cadherin. (A) Cross-section of 3D image data, substrate at bottom. The extracted organoid develops within a lumen. Blue contour indicates the extracted surface. (A’) 3D rendering of the extracted surfaces. (A”) 2D projection. **B-B”** Lightsheet recording of a 16hpf zebrafish embryo, with fluorescently marked nuclei. Anterior left, ventral view. (B) Cross-section of 3D data. (B’) Extracted surfaces with seams of the UV map marked in color. (B”) Corresponding 2D projection with multiple patches for the ventral/dorsal sides of the optic vesicles and the lateral sides of the neural tube. **C-C”** Confocal *z*-stack of *Arabidopsis thaliana* flower buds with a membrane marker. Cross-section of 3D image data. (C’) Extracted surface. (C”) 2D projection (used for cell segmentation in Fig. S1).

### 3 btc can process diverse and complex shapes

We now demonstrate the applicability of `btc` to complex shapes from a range of biological contexts (Fig. 4, data sources in Materials & Methods). Fig. 4A-C shows a confocal *z*-stack of a human neural tube organoid grown on a micropatterned substrate [18]. Tissue cartography offers clear advantages over maximum *z*-projections and as height-map-based approaches like `LocalZProjector` [10], both of which are unable to follow the organoid’s curved apical surface due to overhangs. `btc` supports multichannel data (here, neural and epithelial cadherin).

Next, we extracted the anterior neural tube and developing optic vesicles [19] from an *in-toto* light-sheet recording of a 16hpf zebrafish embryo (Fig. 4A’-C’). This complex shape is beyond the reach of the `ImSAnE` or `TubULAR` software, but can be easily unwrapped along seams following morphological landmarks, selected graphically within `Blender` (Fig. 4B’). Segmenting out this shape from 3D data (in particular, separating it from the surrounding EVL tissue layer), is also greatly streamlined by Blender’s sculpting and modeling tools. The last example are *Arapidospis thaliana* flower buds (Fig. 4A”-B”).

### Quantitative analysis in cartographic projections with the btc Python library

Cartographic projections are not only useful for visualization but also for quantitative analyses: image analysis in 2D is significantly easier than in 3D. However, like a map of the globe, cartographic projections introduce distortion. The `btc` Python library contains a set of tools for mathematically correcting cartographic distortion and mapping points in the projection back into 3D. Tools for shape analysis are also included.

Fig. S1 presents two examples. Fig. S1A-B shows a cell segmentation of the *Arabidopsis* flower buds from Fig. 4C, and uses `btc` to correctly measure cell areas, correcting for cartographic distortion. Fig. S1C takes a closer look at the zebrafish optic vesicle (Fig. 4B). We segmented the cells on the dorsal side of the cup (which folds to form the eye cavity) to carry out *mechanical inference* [9] (Fig. 4D). This method infers mechanical tension from cell shapes. We segmented cells in 2D, and then mapped back to 3D to correctly measure the cell geometry. Under the assumption that mechanical stresses are concentrated along cell-cell interfaces, the interfacial tension can be inferred from the geometry of cell interfaces (Fig. 4D, bottom inset). We find no clear pattern at this developmental stage (Fig. S1D’); in particular, there is no tension cable around the margin of the optic cup (i.e., no “purse string” to drive folding).

## Discussion

Here, we presented `blender tissue cartography` (`btc`), a software suite for analyzing dynamic 3D microscopy data with the 3D editor `Blender`. Tissue cartography extracts and cartographically projects surface-like structures from volumetric data, reducing 3D to 2D data analysis and enabling quantitative insights into animal and plant development. `btc`’s graphical user interface allows iterative design of cartographic projections for diverse and complicated geometries. Via a modular design and by re-purposing high-quality computer graphics software (with a large user base), we standardize tissue cartography workflows and drastically lower barriers to entry for non-experts. We demonstrated `btc` on static and dynamic datasets with complex geometries from four different model organisms, notably on *in-toto* recordings of the zebrafish embryo. The round yolk mass of the zebrafish embryo has made visualization of its anteroposterior axis in its entirety challenging. Typically, embryos must be fixed, dissociated from their yolk, and flat-mounted to provide a full view of the embryonic body [20]. The cartographic projections shown in Fig. 2 make this feasible in a living embryo, enabling unhindered, dynamic observation of somitogenesis and neurulation.

For dynamic data and comparison across samples, registering multiple images to a common reference frame is essential [5]. This task is greatly facilitated by cartographic projections. Using Blender’s UV editor, the user can map anatomically salient features (like the anterior pole of the zebrafish) to defined cartographic positions. Our pipeline for dynamic data allows the user to define the cartographic projection for a reference time point, from which it is algorithmically transferred to subsequent time points. This approach is applicable without modification across a wide range of datasets.

In our experience, `btc` has proven to be user-friendly and easy to learn, with second-year undergraduates and scientists with no computer programming experience able to analyze the data of their interest. `btc` has already been used in publications and preprints [21, 22, 23]. We believe our tool will make the analysis of dynamic 3D surfaces accessible to a broad audience and thus further our understanding of morphogenesis. Approaching development and biological shape via cartographic transformations was originally proposed by D’Arcy Thompson in 1917, and we hope that our work will help connect this vision with the striking microscopy data of the 21^st^ century.

## Methods

### btc add-on and Python library

The `btc` Python library is built using the standard Python scientific computing software stack [24, 25, 26] and uses a lightweight mesh data representation inspired by the `.obj` mesh file format. The geometry-processing library `igl` is used as the main geometry backend [27]. `MeshLab` [16] is used for certain advanced (re)meshing operations. The source code is available on GitHub. The `Blender` add-on is based on the `btc` library code with adaptations to interface with the `Blender` API and is also available on GitHub.

## Data sources

The *Drosophila* blastoderm in Fig. 1A shows a multiview light-sheet microscope recording of a *+;+;H2A:RFP* embryo at stage 6 from Ref. [5]. Fig. 3C-D shows the embryonic *Drosophila* midgut extracted from a multiview lightsheet microscope recording of a *Hand:GFP;+;Hist:GFP* embryo at stage 15 from Ref. [2]. Fig. 4A-C shows a confocal microscope recording of a human neural tube organoid stained for N-cadherin and E-cadherin at 48 hours post BMP addition (organoid protocol initiation) from Ref. [18]. Fig. 4A”-C” shows a confocal recording of *Arabidopsis thaliana* flower buds with plasma membrane marker *pUBQ10::29-1-GFP* [28], shared Dr. An Yan. Permission to display the data was obtained from the respective authors of the respective publications. The zebrafish datasets in Figs. 2, 4B, and S1C were recorded for this manuscript (see next section).

### Zebrafish microscopy

Adult zebrafish of the AB/TU background were used for experiments and housed and bred using standard conditions [29]. All experiments were performed with institutional approval from the University of California, Santa Barbara. Zebrafish embryos were mounted in agarose and imaged in a multi-view light-sheet microscope [30], as described previously [31]. Multiview data was registered and fused as described previously [31, 32] to obtain volumetric image data with isotropic resolution of 0.26*µ*m. We used the following fluorescent reporter lines: *Tg(tbx16:RFP)* [17], microinjected with H2B-RFP mRNA (Fig. 2), *Tg(h2afva:h2afva-GFP)* (Fig. 4), and *Tg(h2afva:h2afva-GFP)kca66; Tg(actb2:mem-Cherry)hm29* [33] (Fig. S1). For mRNA injection, H2B-RFP mRNA (a gift from the Wallingford lab) was synthesized from NotI-linearized pCS2+-H2B-RFP plasmid using the SP6 mMessage mMachine kit (Invitrogen). A total of 50 pg of H2B-RFP was microinjected into single-cell embryos with pulled glass needles.

### Data processing

Data was processed following the pipeline described in the main text methods section. In brief, all volumetric datasets were segmented using `ilastik` with additional post-processing via level set methods for the zebrafish dataset. The volumetric segmentations were converted to meshes using the `btc Blender` add-on, and remeshed for improved mesh quality where necessary using either `Blender` or `MeshLab`. For the embryonic *Drosophila* midgut data, we used surface meshes generated in Ref. [2]. UV maps were generated using the “Minimum Stretch” and “Angle Based Flattening” algorithms in `Blender` [12] after graphical seam placement. To obtain a cylinder-like projection for the embryonic midgut data, we used the “Pin” and “Align” tools of the UV editor to ensure straight seams. All 2D projections and 3D renderings were created in `Blender`, using the `btc` add-on. Cell segmentations were carried out using Tissue Analyzer [34], and mechanical inference was carried out following the method of Ref. [9] and using the codebase Ref. [35].

## Acknowledgments

We thank Noah Mitchell, Pieter Derksen, Fridtjof Brauns, An Yan, Dillon Cislo, and all members of the Streichan lab for helpful discussions, providing sample data, and software testing. NHC was supported by NIGMS R35-GM138203, NSF PHY:2210612, and a PCTS fellowship.

## Author contributions

NHC designed and developed the software, analyzed the data, and wrote the manuscript. CR provided expertise in 3D modeling and the `Blender` software and created visualizations. SW recorded the zebrafish data. MFL recorded the *Drosophila* data. SJS supervised the project.

## Supplementary Information

### A Computational representation of surfaces and projections

We represent surfaces as triangular meshes with vertices and triangles (𝒱, 𝒯), saved as standard `.obj` files (several previous tissue cartography tools use custom data formats, complicating interoperability with other software). We use Poisson reconstruction or the marching cubes algorithm to construct these meshes from binary segmentations.

UV maps of a surface ((𝒱, 𝒯) are represented in a standardized fashion as a set of 2D UV vertices in the unit square (𝒱_uv_ ⊂ [0, 1]^2^, and a set of UV faces 𝒯_uv_ that are in one-to-one correspondence with the 3D faces. Therefore, cartographic maps and projections always lie in the unit square. The mesh and its UV map can be exported to or loaded from an `.obj` mesh.

Given the SOI mesh with a UV map, `btc` uses an interpolation algorithm to project voxel intensities from the volumetric data onto the 2D UV plane. The interpolation scheme is designed so that the resolution of the 2D projection does not depend on that of the mesh, i.e., a coarse, lightweight mesh can still be used for high-resolution projections. `btc` features a second, simpler projection algorithm that evaluates 3D image intensities at mesh vertices. This vertex-shader algorithm does not require a UV map and can be recomputed rapidly.

### B Vector calculus on triangular surfaces

We also provide a simple implementation of vector calculus on curved surfaces, for instance, to analyze morphogenetic tissue flows obtained from particle image velocimetry (PIV) or cell tracking in 2D projections [4, 2]. In brief, vector and tensor fields on a surface are represented by their Cartesian 3D coordinates and then separated into tangential and normal components. Using standard finite-element operators on the mesh, gradients can be calculated componentwise and combined to form the curved-surface generalizations of familiar operators like div, grad, and curl [36], and approach accessible to non-experts in differential geometry. These methods draw on tools from “discrete differential geometry”, developed in computer graphics [37].

### C Batch processing and dynamic datasets

Often, one is interested in dynamic datasets (movies), or multiple recordings of similarly shaped objects (e.g., multiple *Drosophila* blastoderm stained with different markers[5]). In `btc`, timelapse datasets are represented frame-by-frame (one surface mesh and volumetric image file per timepoint). Such collections of SOIs can be batch-processed via the `btc` add-on or Python library. It is highly desirable to use “the same” UV map for each time point, both to facilitate comparison of the 2D projection across time points or samples and because it would be cumbersome to manually define a UV map each time. There are two possible strategies.

The first possibility is to compute a UV map for each mesh individually, but using a consistent algorithmic procedure. This approach works well for simple shapes that can be unwrapped by axis, cylindrical, or spherical projections, which can be batch-computed in `Blender`. However, for more complex shapes (in particular, if unwrapping requires seams), choosing or designing a suitable algorithm can require significant computer vision expertise and, crucially, cannot be done graphically and interactively.

Instead, in `btc`, the user defines a UV map for a selected *reference mesh* – e.g., the first or last frame of a movie, or an “idealized” version of the imaged shapes – using `Blender`’s graphical UV editor. The reference mesh is then algorithmically mapped onto the meshes of the remaining time points (the *target meshes*), a process that we refer to as *surface-to-surface alignment* (Fig. 3A). This allows *transferring the UV map* from the reference mesh to all other meshes and hence creates coherent 2D projections across time points or samples. In particular, the layout of the 2D projection in the UV square does not change. For the case of multiple samples, surface-to-surface alignment allows combining recordings of e.g. different markers into a single “atlas” [5]. Surface-to-surface registration has received significant attention in the computer graphics community. In `btc`, we implement three algorithms, all based purely on surface geometry (i.e., they make no use of the 3D image data):

1. **Rigid-body alignment**. The reference mesh is translated, rotated, and scaled to match the target mesh (Fig. 3B).
2. **Closest-point projection (shrink-wrapping)**. The reference mesh is first rigidly aligned to the target, and then each vertex of the reference mesh is projected to the closest point on the target surface (Fig. 3B’). This operation is combined with smoothing to remove “creases”.
3. **Moebius alignment [38]**. Both reference and target mesh are mapped to a reference shape (a disk, a punctured disk, or a sphere, depending on the surface topology) in a way that preserves triangle angles. Mathematically, this *conformal map* to the reference shape is guaranteed to be almost unique (technically, `btc` uses harmonic maps [39].). The remaining freedom lies in carrying out a *Moebius transformation* (e.g., a rotation). The Moebius transformation is chosen to best align the geometry of the reference and target mesh (Fig. 3B”).

**Figure S1:**
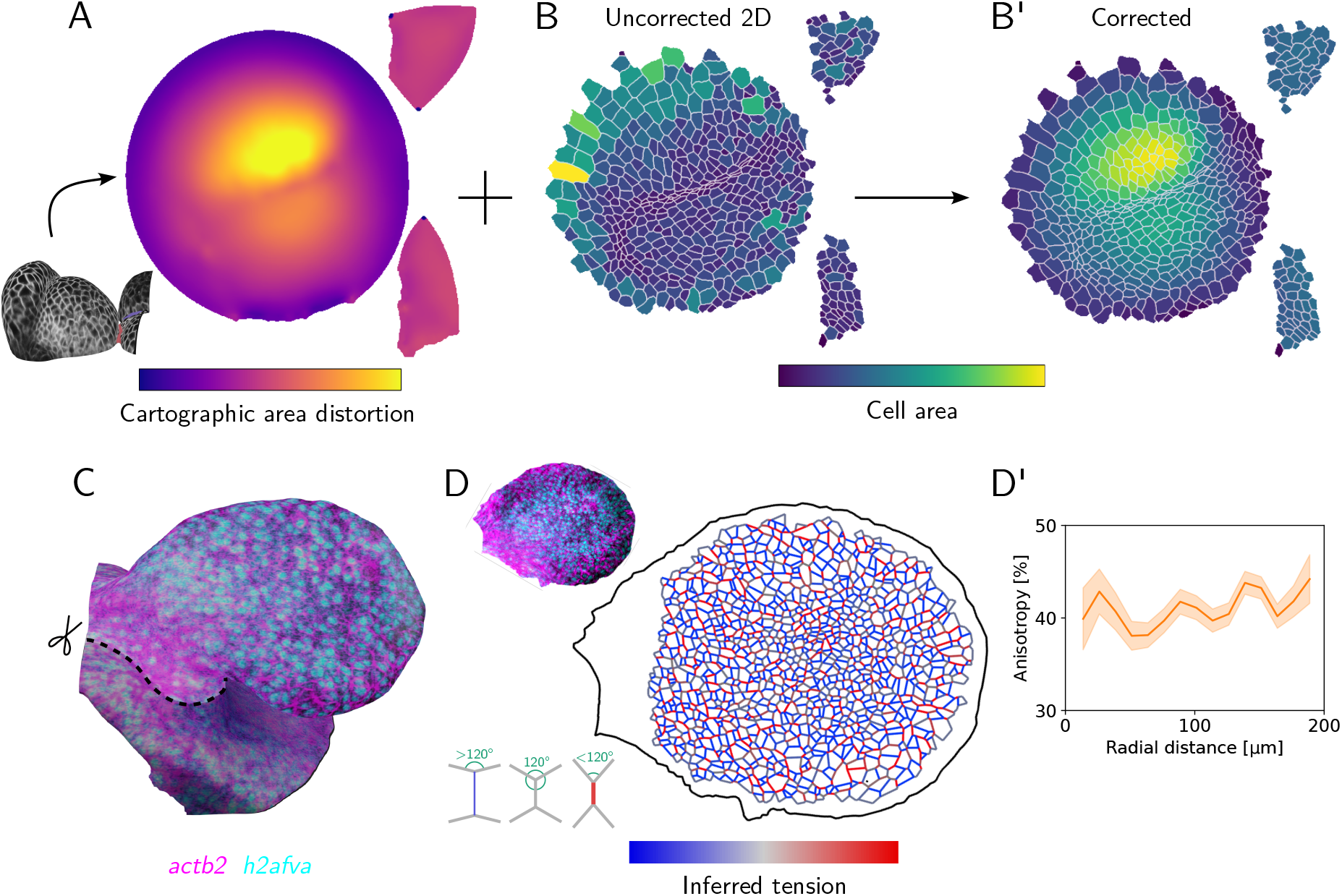
Quantitative analysis of data on curved surfaces with the btc Python library. **A** Area distortion of UV map for the *Arabidopsis thaliana* data (inset, Fig. 4C”). **BB’**(D’-D”) Segmentation of 2D projections, colored by (B) 2D cell areas and (B’) cartographically corrected, 3D cell areas. **C** 3D mesh of the developing optic zebrafish optic cup at 16hpf (see Fig. 4B), dorsal-anterior view (cyan: h2afva (nuclei), magenta: actb2 (membrane)). Attachment to the neural tube is visible on the left. Analyzed portion of the surface in bright. **D-D’** Analysis of cell mechanics on the apical surface of the optic cup. Projected nuclear and membrane signal (top inset) was used to segment out cells and infer relative interfacial tensions on cellcell interfaces from cell geometry (bottom inset). (D’) Tension anisotropy (deviatoric part of tricellular vertex stress tensor [9]) shows no clear center-to-margin pattern, indicating the absence of a tension cable around the optic cup margin.

For dynamic datasets, surface-to-surface registration can be done iteratively (i.e., first map reference to the first frame, then the result to the second frame, etc.), significantly reducing its difficulty (Fig. 3C).

Thanks to `btc`’s extensible Python interface, external libraries can also be used. The above algorithms are fully automatic and do not require the user to specify point-to-point correspondences between reference and target mesh, which is time-consuming and can introduce bias in the absence of clear shapes enabling interest point placement. The Moebius algorithm is highly robust and will find a surface-to-surface map even between two strongly deformed meshes [38].

